# Vascular physiology drives functional brain networks

**DOI:** 10.1101/475491

**Authors:** Molly G. Bright, Joseph R. Whittaker, Ian D. Driver, Kevin Murphy

## Abstract

We present the first evidence for vascular regulation driving fMRI signals in specific functional brain networks. Using concurrent neuronal and vascular stimuli, we collected 30 BOLD fMRI datasets in 10 healthy individuals: a working memory task, flashing checkerboard stimulus, and CO_2_ inhalation challenge were delivered in concurrent but orthogonal paradigms. The resulting imaging data were averaged together and decomposed using independent component analysis, and three “neuronal networks” were identified as demonstrating maximum temporal correlation with the neuronal stimulus paradigms: Default Mode Network, Task Positive Network, and Visual Network. For each of these, we observed a second network component with high spatial overlap. Using dual regression in the original 30 datasets, we extracted the time-series associated with these network pairs and calculated the percent of variance explained by the neuronal or vascular stimuli using a normalized R^2^ parameter. In each pairing, one network was dominated by the appropriate neuronal stimulus, and the other was dominated by the vascular stimulus as represented by the end-tidal CO_2_ time-series recorded in each scan. We acquired a second dataset in 8 of the original participants, where no CO_2_ challenge was delivered and CO_2_ levels fluctuated naturally with breathing variations. Although splitting of functional networks was not robust in these data, performing dual regression with the network maps from the original analysis in this new dataset successfully replicated our observations. Thus, in addition to responding to localized metabolic changes, the brain’s vasculature may be regulated in a coordinated manner that mimics (and potentially supports) specific functional brain networks. Multi-modal imaging and advances in fMRI acquisition and analysis could facilitate further study of the dual nature of functional brain networks. It will be critical to understand network-specific vascular function, and the behavior of a coupled *vascular-neural network*, in future studies of brain pathology.

## INTRODUCTION

Imaging neuroscience has advanced a new theory of brain function based on the interconnectedness of neuronal activity in multiple brain regions^1^. These regions form structural and functional networks that are consistent across individuals^2^ and intrinsic to brain activity during active processing or in the resting state^1,3^. To provide efficient and targeted support for such neuronal networks, we hypothesize that the cerebrovasculature has also evolved characteristics of functional networks.

It is well established that local blood flow is tightly coupled to local neuronal activity to protect brain metabolism^2,4^. This coupling is what underpins the Blood Oxygen Level Dependent contrast mechanism in functional magnetic resonance imaging (fMRI) of brain activity, and this technique has been used to characterize functional networks in thousands of neuroimaging studies of the human brain^5^. Several functional brain networks are robustly identified in human subjects, in both task-activation and resting-state datasets^2,3,6^, and are frequently characterized in patient cohorts to better understand the mechanisms of pathology.

However, the vasculature can also regulate local blood flow in response to physical and chemical signals, independently of local neuronal activity^7^. Inhalation of air with elevated levels of carbon dioxide (CO_2_, a potent vasodilator) is frequently used to drive a vascular response and a resulting BOLD signal increase. The resulting maps of *cerebrovascular reactivity* are frequently used to assess impairment in vascular function, such as in multiple sclerosis^8^, Alzheimer’s disease^9^, and stroke^10,11^. This type of hypercapnia challenge impacts arterial blood gas tensions systemically, influencing all of the cerebrovasculature simultaneously. However, even in healthy individuals the characteristics of local vascular responses to CO_2_ vary across the brain^12^ and may demonstrate regions of coordinated vascular regulation.

In a previous fMRI study to improve our methodology for mapping this regional variation in vascular regulation, we used a breath-hold paradigm to induce changes in end-tidal CO_2_^13^. In further exploratory analyses, we averaged the resulting BOLD-weighted data across subjects and used Independent Component Analysis (ICA) to decompose the data into spatially independent “network” maps and associated time-series. In these results, we identified that the Default Mode Network (DMN) was represented by two components: one DMN time-series showed clear BOLD signal increases lagging the end-tidal CO_2_ effects, thereby reflecting the vasodilatory effect of the stimulus. Interestingly, the second DMN component time-series exhibited BOLD signal *decreases* during and preceding the actual breath-hold itself, potentially reflecting deactivation of this network and reduced neural activity during the active, attentional portion of the paradigm. (See **Supplemental Figure 1** for a summary of these preliminary results.)

Based on this observation, we hypothesize that functional brain networks may be comprised of two, distinct but coupled systems: one primarily driven by neuronal activity and one driven more by vascular regulation. Extending this premise, vascular regulation may occur in a coordinated manner across multiple, long-distance brain regions, mimicking or contributing to known functional brain networks.

To test this hypothesis, we developed a protocol to probe both physiological systems across multiple brain networks, employing concurrent and orthogonal neuronal and vascular stimuli. We decompose the resulting BOLD signal changes using ICA and identify the relative influence of neuronal and vascular factors on functional brain networks. Our results provide further evidence for the dual-nature of functional brain networks, and highlight the importance of characterizing *vascular* function as well as neuronal function within specific brain networks.

## METHODS

Whole-brain functional MRI neuroimaging data were collected during stimuli designed to simultaneously probe neuronal and vascular systems throughout the brain. These data were then decomposed to identify network structures reflecting either neuronal or vascular mechanisms.

### Neuronal and vascular stimuli

A 3-back working memory task (centrally presented, digits 0-9, presented for periods of 0.5 s at 1.5 s intervals) was delivered in a 30-second block design, with an extended (60 s) off-period in the middle of the paradigm. Participants were asked to press a button when the digit presented was the same as that presented three stimuli previously. A visual stimulus consisting of a radial flashing checkerboard pattern was also presented in a block design in the second half of the scan (8 Hz, 70% contrast, with neutral center to allow simultaneous presentation of the working memory task). These stimuli were presented using a rear projector and screen viewed through a mirror on the head coil.

During these neuronal tasks, four 1-minute blocks of passive hypercapnia were used as a concurrent vascular stimulus. A gas mixture with increased levels of carbon dioxide (CO_2_) was delivered to the subject via a face mask, manually adjusted to target an end-tidal CO_2_ increase of +5 mmHg. Inhalation of CO_2_ alters arterial blood gas tensions, which results in vasodilation and enhanced blood flow throughout the body. It is known that the response of local vessels to this systemic stimulus varies across the brain, in amplitude and dynamics, and these variations can be observed using BOLD fMRI^12^. We hypothesize that these variations in the response to hypercapnia will reveal spatial patterns of coordinated vascular regulation, or vascular networks.

All three stimulus paradigms were designed to be mutually orthogonal. A schematic of each stimulus is presented in **Figure 1**. The neuronal stimulus timings were convolved with a standard gamma variate hemodynamic response function. The end-tidal CO_2_ data were extracted using bespoke code (MATLAB, MathWorks, Natick, MA, USA), convolved with the same hemodynamic response function, and used as a scan-specific measure of the vascular stimulus evoked by the hypercapnia challenge^13^.

**Figure 1.**
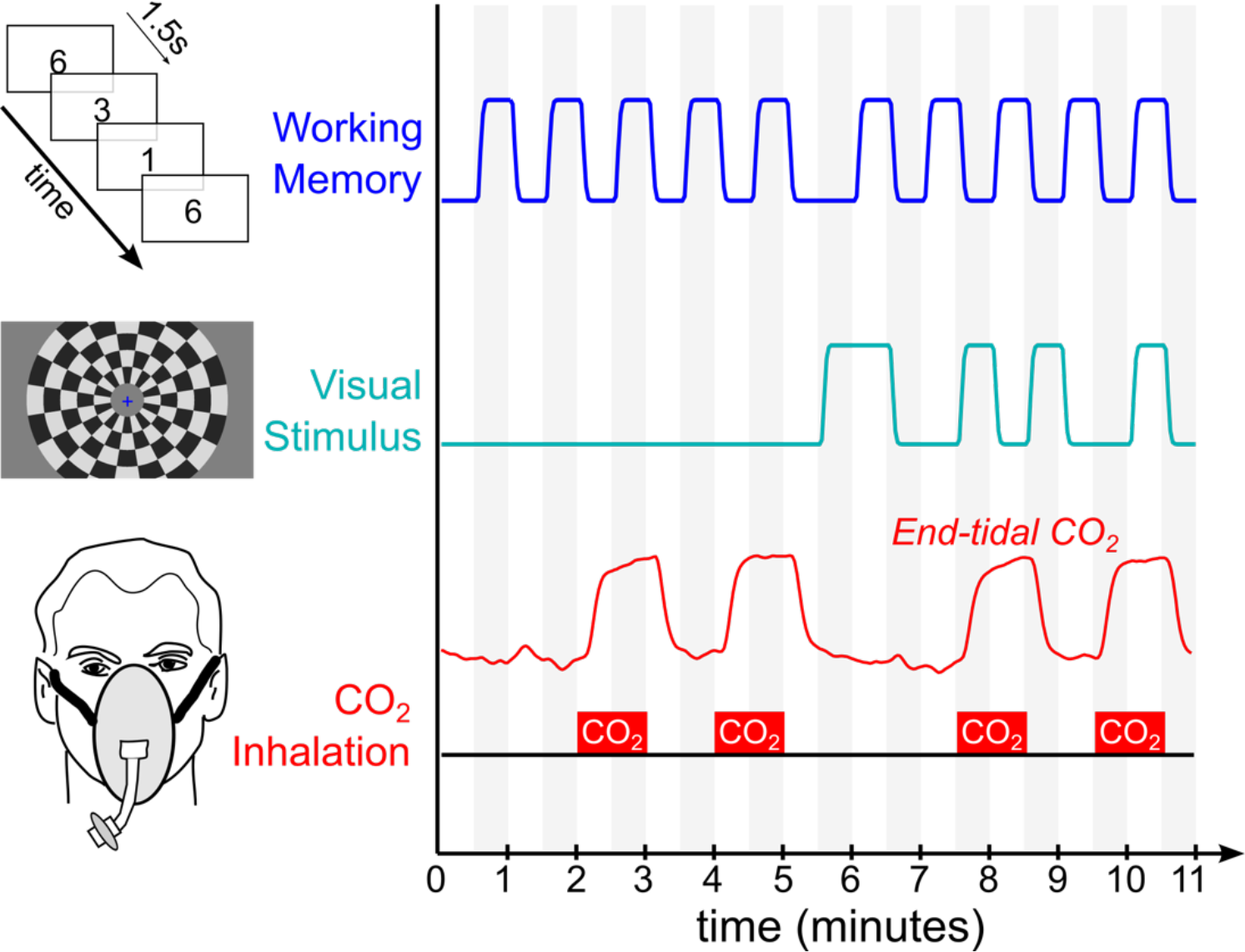
Schematic of neurovascular stimulus paradigm. The neuronal stimuli (working memory task and flashing checkerboard pattern) were presented in a block design, which was convolved with a hemodynamic response function to model the resulting BOLD signal. Four 1-minute blocks of hypercapnia were induced via gas inhalation, and modeled in a subject-specific manner by extracting the end-tidal CO_2_ data and convolving with a hemodynamic response function.

In a follow-up study (*Replication*) that did not involve the hypercapnia stimulus, the participant’s end-tidal CO_2_ levels were allowed to fluctuate naturally, and a nasal cannula was used to monitor respiratory gas content in lieu of the face mask.

### Data acquisition

Ten healthy subjects (aged 30 ± 5 years, 3 female) were scanned using a 3T GE HDx scanner (Milwaukee, WI, USA) equipped with an 8-channel receive head coil. Three identical functional task scans, each lasting 11 minutes, were acquired using a BOLD-weighted gradient-echo echo-planar imaging sequence (TR/TE=2000/35 ms; flip angle=90°; FOV=22.4 cm^2^; matrix=64x64; 35 slices, slice thickness=4 mm; resolution=3.5×3.5×4.0 mm^3^, 5 dummy scans followed by 330 volume acquisitions), for a total of 30 datasets of 11 minutes duration each. A whole-brain high-resolution T1-weighted structural image was acquired (resolution=1.0×1.0×1.0 mm^3^), for the purpose of image registration, and a fieldmap was acquired (matching functional scan acquisition, echo spacing=0.32 ms) for correction of image distortion artifacts.

Expired gas content was continuously monitored via a sampling port on the face mask, and O_2_ and CO_2_ data were recorded using a capnograph and oxygen analyzer (AEI Technologies, PA, USA). End-tidal CO_2_ data were extracted and convolved with a hemodynamic response function (**Figure 2**); the hypercapnia achieved in this protocol (averaged across the study) was 5.8 ± 1.1 mmHg above baseline levels.

**Figure 2.**
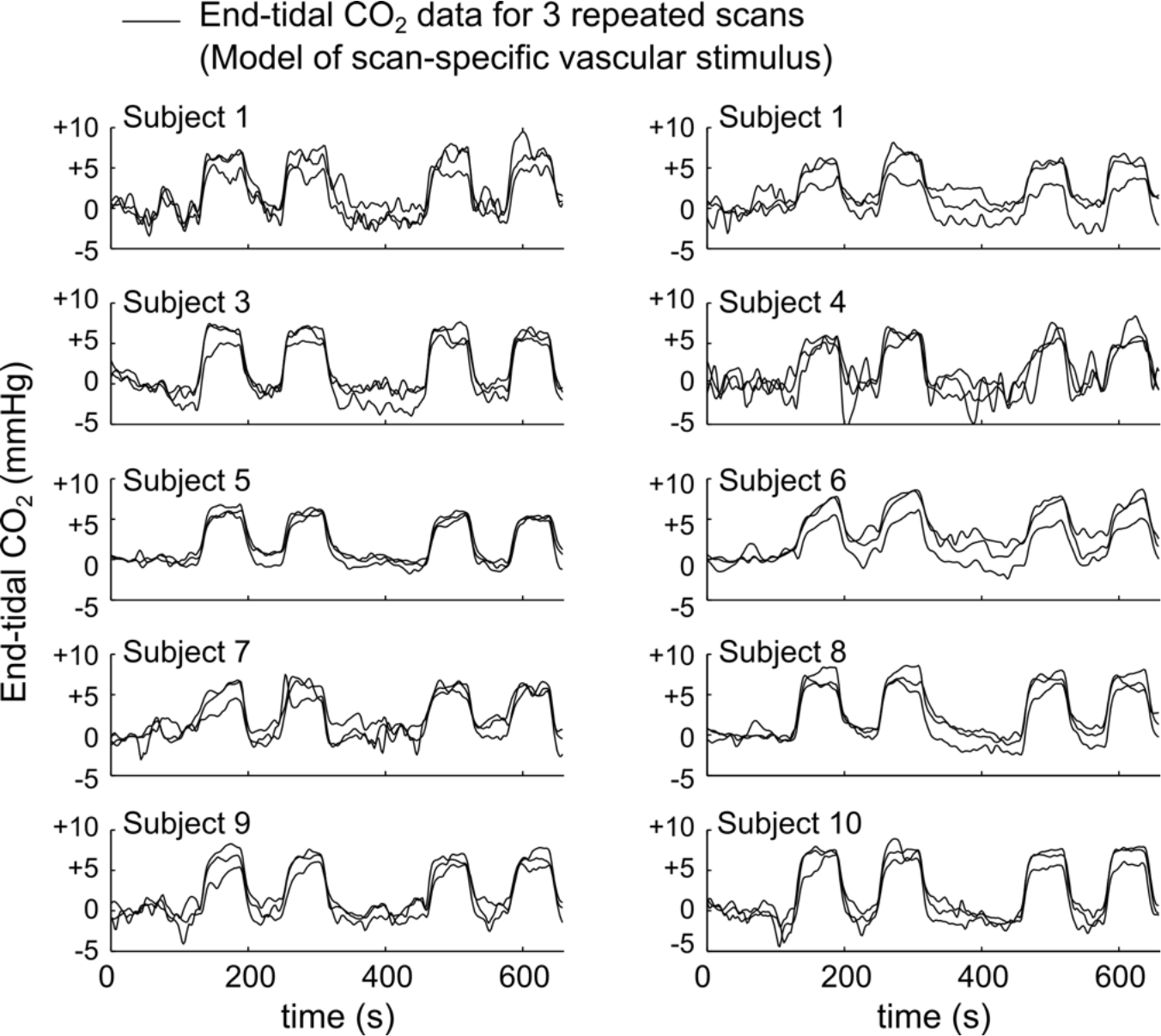
End-tidal CO_2_ data for all scans. Data for 30 scans (3 repeated scans per participant) were convolved with a hemodynamic response function, and represented the scan-specific vascular stimulus. For illustration purposes, data were normalized to the baseline end-tidal CO_2_ level (mean value in the first 100 seconds of the scan).

The study cohort size was not determined via calculation, but was determined to be sufficiently large given the literature studying neuronal^2^ and vascular^14^ networks. This study was approved by the Cardiff University School of Psychology Ethics Committee, and all volunteers gave written informed consent.

### fMRI data processing

Following motion correction (AFNI^15^, http://afni.nimh.nih.gov/afni), brain extraction (BET, FSL^16^), distortion correction and slice timing correction (FEAT, FMRIB’s Software Library, Oxford, UK^17^) all functional datasets were registered to the corresponding high-resolution T1-weighted image, which was in turn normalized to the MNI-152 brain template (MNI152, nonlinearly derived, McConnell Brain Imaging Centre, Montreal Neurological Institute, McGill University, Montreal, Quebec, Canada) using FMRIB’s Non-linear Image Registration Tool (FNIRT, FSL^18^).

Datasets were then detrended using second order polynomials and converted into units of percentage change (%BOLD). The 30 pre-processed %BOLD datasets were averaged together to reduce the influence of any signals not time-locked to the stimulus paradigm (i.e., resting fluctuations and other noise confounds).

### Network analysis

The average dataset was decomposed into spatially independent networks using independent component analysis, as implemented in the MELODIC tool in FSL (dimensionality fixed to output 30 components; each comprised of a network map and associated time-series). Because BOLD-weighted signals are both directly influenced by changes in the vasculature, and indirectly influenced by neuronal activity via neurovascular coupling, the temporal characteristics of signal changes were used to determine whether they reflect primarily neuronal or vascular mechanisms. Three ‘neural networks’ were identified using temporal correlation values of the component time-series and stimulus timings as follows. The component with the maximal negative correlation with the 3-back stimulus was identified as the neuronal Default Mode Network (DMN), which is robustly de-activated during working memory tasks^19–21^. The component demonstrating maximum positive correlation with the 3-back stimulus was identified as the neuronal Task Positive Network (TPN)^22,23^. Finally, the component exhibiting maximum positive correlation with the visual stimulus was identified as the neuronal Visual Network (VN).

Component maps were thresholded using a mixture model and an alternative hypothesis testing approach^24^ (threshold level 0.5) and spatial similarity between all 30 components was quantified using Dice’s overlap coefficient^25^. For each neuronal network, we identified the additional component map with the greatest spatial overlap. Thus, three pairs of spatially coupled components were identified.

Finally, dual regression^26^ was used to extract the time-series associated with these 6 components within each of the original 30 datasets. The normalized R^2^ value was defined as the time-series variance (R^2^) explained by one stimulus normalized by the variance explained by the full stimulus model, which is the percentage of explained variance attributed to one stimulus. Paired two-tailed Student t-tests were used to compare the normalized R^2^ values of each component pairing (DMN, TPN and VN) for each of the stimuli. Normality of the pair-wise differences was assessed using the Lilliefors test, and significant non-zero differences in the temporal signatures of the component pairs were identified (*p<0.05, Bonferroni corrected for multiple comparisons).

### Replication and generalizability

Eight of the original 10 subjects were scanned, three times each, using a reduced stimulus paradigm consisting of only the working memory and visual stimuli. Thus, in these scans, end-tidal CO_2_ was allowed to fluctuate naturally rather than be driven by the gas inhalation stimulus. As such, these scans follow more “typical” task-activation fMRI experiments, and they will allow us to assess the replicability and generalizability of our observations.

Data were pre-processed as described above. Because the end-tidal CO_2_ levels were allowed to fluctuate naturally, these fluctuations will vary across scans and individuals would not be robustly present in the group-average dataset. Thus, independent component analysis could not be applied to isolate vascular-neuronal network pairs in the averaged data as was done in the original analysis. Instead, using dual-regression the time-series associated with the network maps identified in the *original* dataset were extracted for the *replication* data.

## RESULTS

Following Independent Component Analysis, three resulting components were identified as “neuronal networks”, demonstrating maximum temporal correlation with the neuronal stimulus paradigms: the Default Mode Network, Task Positive Network, and Visual Network were identified (**Figure 3**).

**Figure 3.**
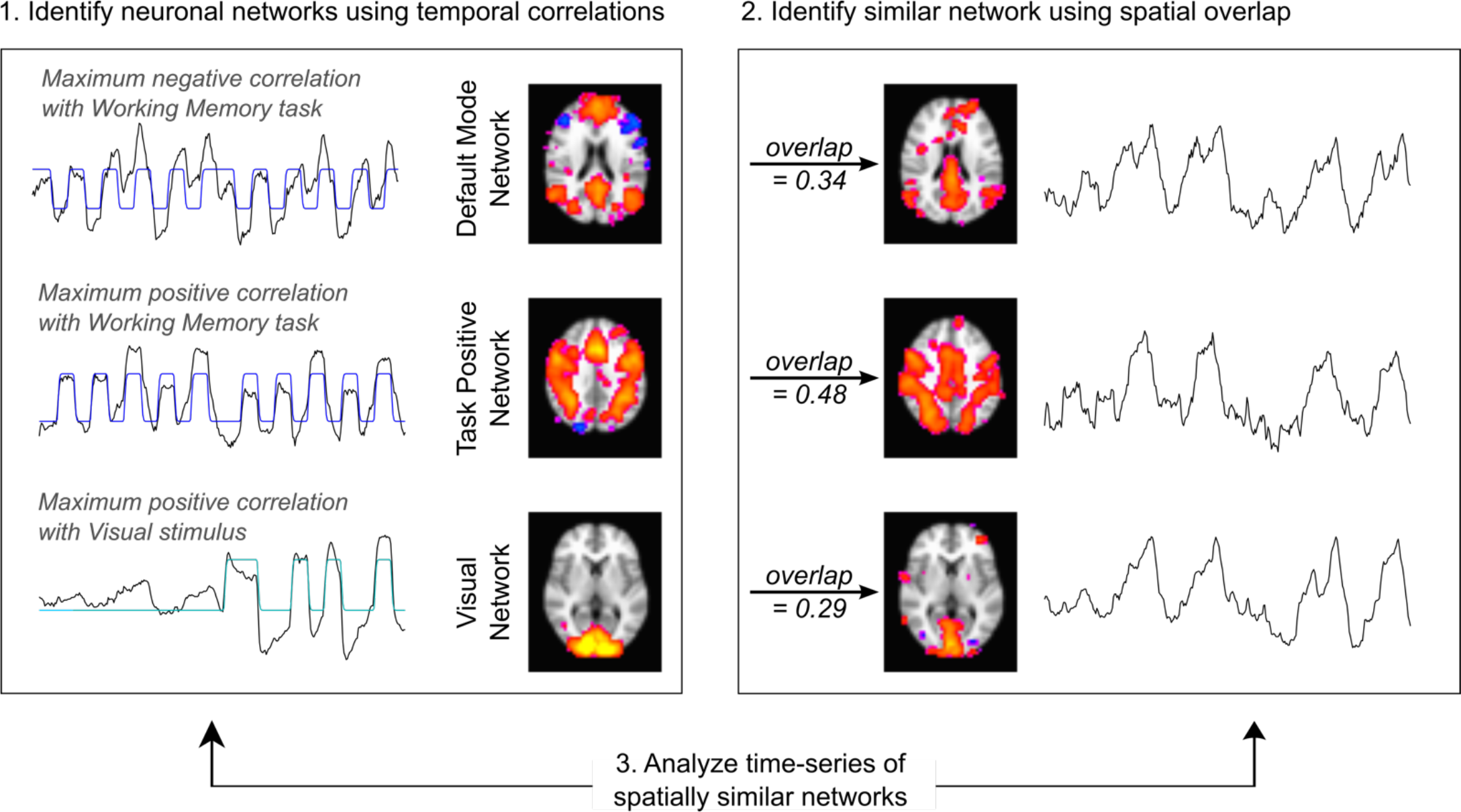
Identification of spatially similar component pairs for three functional brain networks. 1) The three components with maximum temporal correlation with the neuronal stimuli were identified as ‘neuronal’ networks. 2) For each neuronal network, an additional component with the maximal spatial overlap was identified. 3) The temporal characteristics of these spatially similar components were used to assess the underlying neuronal or vascular mechanisms.

Using dual-regression, we extracted the time-series associated with each of these components in the original datasets (all time-series provided in **Supplemental Figure 2**). Each time-series was analyzed to determine the relative contributions of the neuronal and vascular stimuli to the BOLD contrast dynamics, as summarized by the normalized R^2^ values described above. We observed that all three functional brain networks probed in our study were composed of spatially similar pairs of components where one was significantly more associated with the appropriate neuronal stimulus and the other significantly more associated with the vascular stimulus (**Figure 4**).

**Figure 4.**
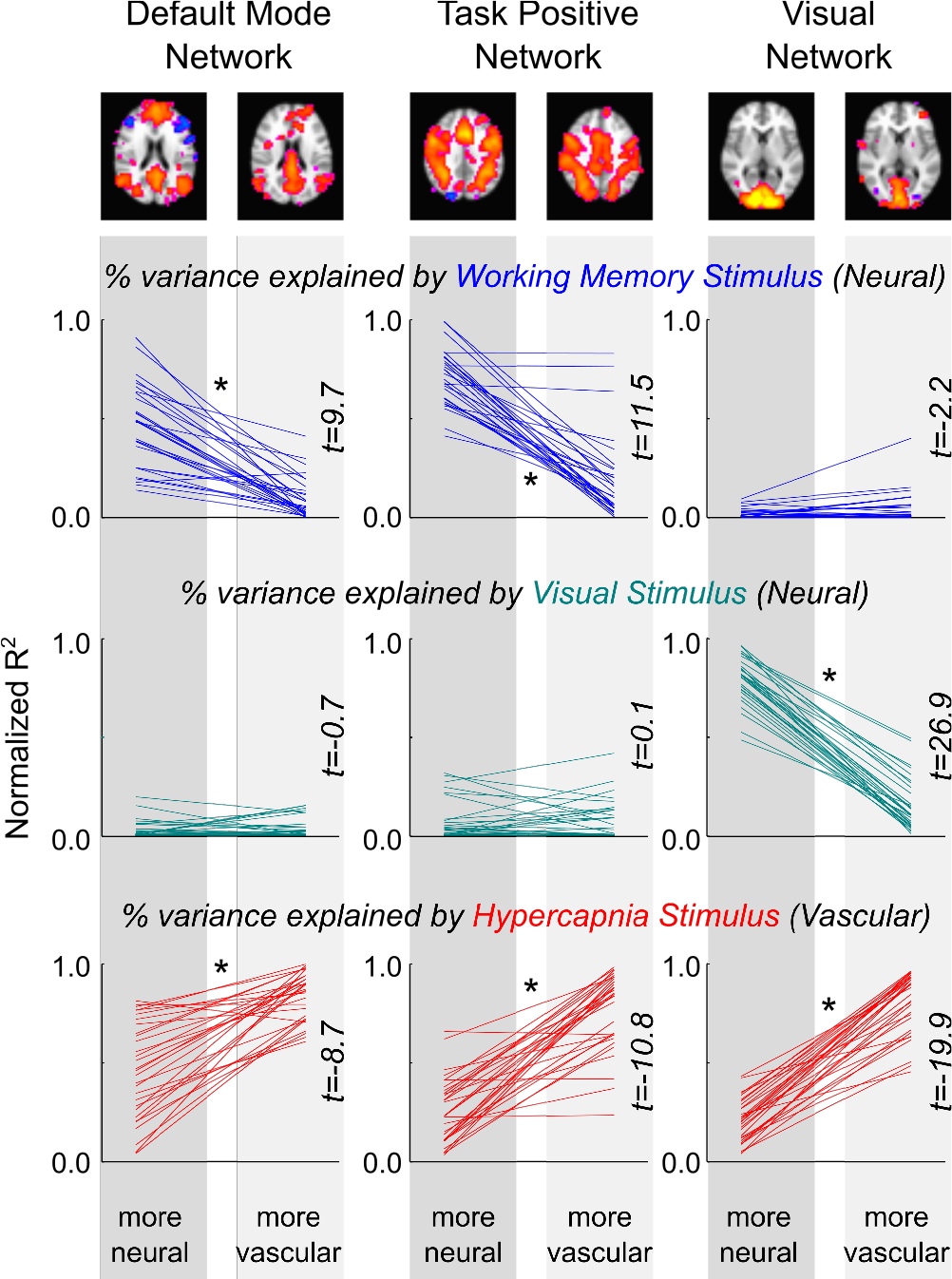
Neuronal and vascular contributions to spatially similar network components. Using dualregression, the component time-series for the three network pairs (Default Mode Network, Task Positive Network, Visual Network) were obtained in the original 30 datasets. The normalized R^2^ values (percentage of explained variance) were calculated for each stimulus. For each functional brain network pair, one was found to be significantly more associated with the appropriate neuronal stimulus and the other significantly more associated with the vascular CO_2_ stimulus (*p<0.05, paired t-tests, corrected for multiple comparisons).

In Fig. 4, the networks on the left of each pairing were identified as maximally temporally correlated with the neuronal stimuli; thus, by design, the working memory stimulus and visual stimulus explain a large proportion of the signal variance (high normalized R^2^ values). Interestingly, the networks on the right of each pairing, which were identified as being spatially similar, show a significantly reduced relationship with these neuronal stimuli. The bottom row shows that all networks show a relationship with the hypercapnia stimulus (represented by the end-tidal CO_2_ data from individual scans), however the networks on the right of each pairing show significantly greater normalized R^2^ values in all cases. Combined, these results suggest that the pairs of spatially similar networks consist of one network representing the neuronal stimuli and one network more reflective of the vascular stimuli. (Note, as a control, we see the expected minimal relationship between the DMN and TPN and the visual stimulus, or the VN and the working memory stimulus.)

When examining the *Replication* dataset, similar phenomena were also observed (**Figure 5**): the normalized R^2^ values demonstrate the same differentiation between the more ‘neuronal’ and more ‘vascular’ components. This demonstrates that our observations are also present in more “typical” fMRI data in the absence of overt hypercapnia challenges, although it is clear that the effects are more variable across individual scans. However, we observe one new effect in the *Replication* data that was not present in the original results. Specifically, the working memory stimulus explains significantly more variance in the “more vascular” VN, whereas no relationship was found in the original data (Fig 5, top right plot).

**Figure 5.**
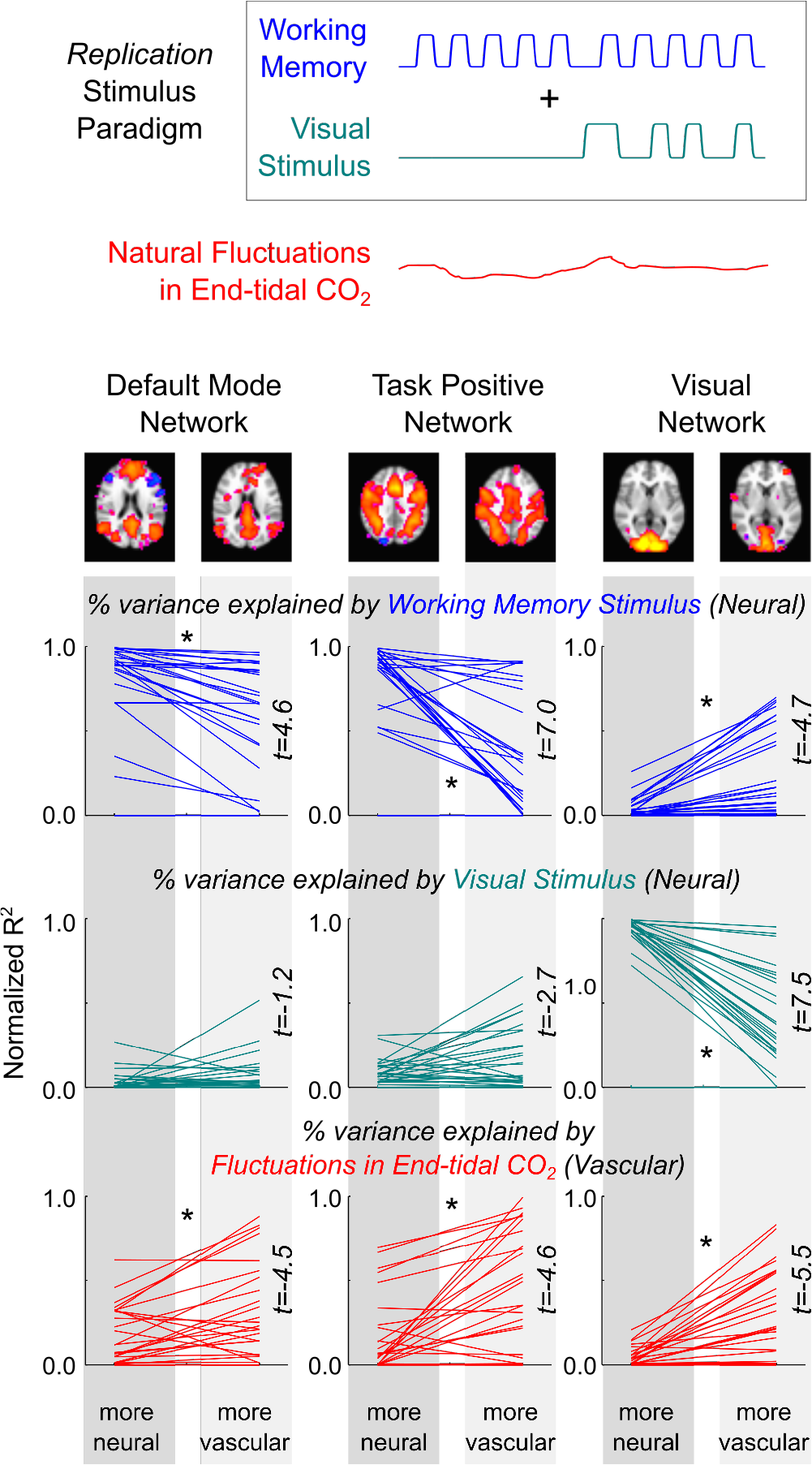
The spatial maps extracted in the original dataset were applied to a second dataset to test the replicability and generalizability of our primary observations. In these new data, no hypercapnia stimulus was administered and end-tidal CO_2_ was allowed to fluctuate naturally. Eight of the original 10 participants were re-scanned, 3 times each, using only the working memory and visual stimuli. The networks identified in the original data were regressed onto the new data, and the associated time-series were extracted and analyzed as before. Significant differences in the normalized R^2^ values, in good agreement with the original observations in the first study, are indicated by asterisks (*p<0.05, paired two-tailed Student t-tests, Bonferroni corrected for multiple comparisons). Note an unexpected, significant relationship between the “more vascular” Visual Network data and the working memory stimulus, not observed in the original dataset (Fig. 4).

Why is the working memory stimulus driving signal fluctuations in the “more vascular” *visual* network component in the *Replication* data? The 3-back task was presented visually, so it is plausible that it would activate the visual processing systems. However, this is not observed in the original dataset, suggesting another mechanism is responsible. It is also known that task-correlated breathing changes are a common, confounding contributor to fMRI data^27^, and this end-tidal CO_2_ may become time-locked to the neural stimulus. Indeed, **Figure 6A** presents the end-tidal CO_2_ time-series of all scans in the *Replication* dataset, showing a strong negative correlation between the group-average end-tidal CO_2_ trace and the working memory stimulus design (Pearson correlation coefficient r=−0.68). Averaging over the ten blocks of the 3-back task, this coupled relationship is even more apparent (r=−0.91). Furthermore, **Figure 6B** shows the BOLD signal changes evoked by the working memory stimulus in the two VNs, clearly showing a BOLD signal *decrease* in the “more vascular” network; this agrees with the concurrent *decrease* in end-tidal CO_2_, while directly countering the argument that the working memory stimulus cues activate the visual cortex (which would be a *positive* BOLD signal change). Thus, the seemingly paradoxical relationship between the “more vascular” VN time-series and the working memory task paradigm is likely caused by vascular physiology becoming time-locked to that neuronal stimulus.

**Figure 6.**
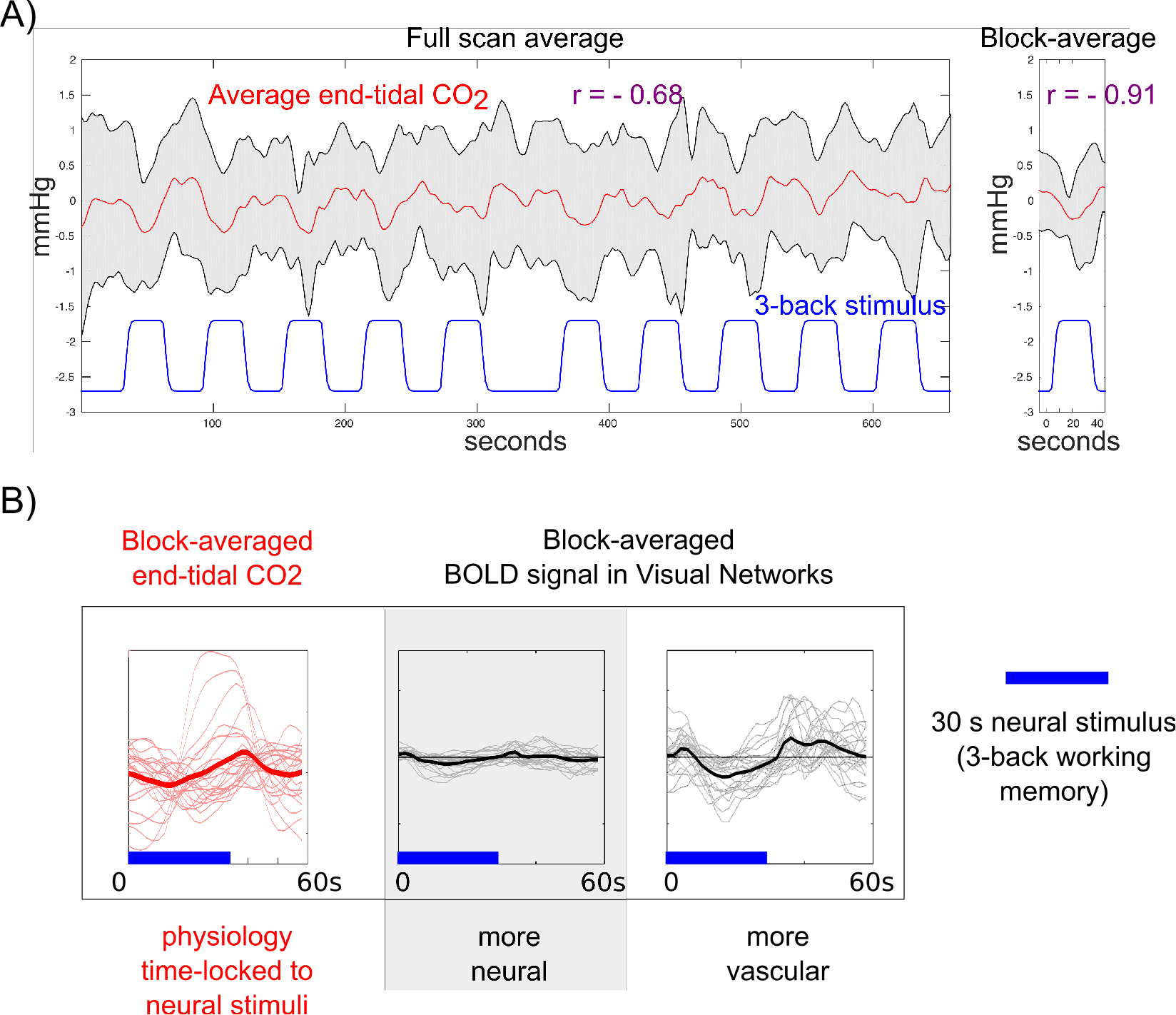
Evidence for task-correlated changes in vascular physiology and its effect on the “more vascular” networks in the *Replication* dataset. **A)** In the absence of a hypercapnia gas inhalation stimulus, end-tidal CO_2_ fluctuated with each individual’s natural variations in breathing. The group average end-tidal CO_2_ trace across all scans in the *Replication* dataset (red, standard deviation shown in gray) is plotted, with the 3-back working memory stimulus paradigm (blue) provided as a reference. The block average across the 10 blocks of the 3-back task is also shown. Pearson correlation coefficients between the CO_2_ data and the stimulus are given (r=−0.68 across the entire time-series, r=−0.91 in the block-average data). **B)** The average end-tidal CO_2_ data and the block-average BOLD response evoked by the 3-back working memory task (blue bars) in the “more neural” and “more vascular” visual networks (thin lines represent the data from each individual scan, thick lines represent the average of 30 scans). These results demonstrate that the task-correlated changes in end-tidal CO_2_ appear to drive the signal fluctuations in the “more visual” network. Because these effects manifest as negative BOLD signal changes time-locked to the working memory task, and visual activation during the working memory task would evoke positive BOLD signal changes, this is further evidence for a vascular driver of this functional network system.

## DISCUSSION

Our findings provide the first evidence for network-specific behavior of cerebrovascular regulation, and suggest the brain’s blood supply may be regulated in networks that spatially mirror known neuronal networks. Using ICA to decompose group-averaged fMRI data, we identified three functional networks associated with working memory and visual stimuli. In the remaining components, three additional networks were identified to have similar spatial features and high spatial overlap as measured by the Dice coefficient. The time-series of these spatially-similar networks were dominated by the vascular stimulus. Although the inhaled carbon dioxide challenge used as a vascular stimulus in this study is known to induce *systemic* vasodilation and BOLD signal increases^28^, our results suggest that the vasodilatory effects show regional variation and that may drive BOLD signal changes in specific functional brain networks or sub-networks.

The spatial similarity of the “more vascular” networks and neuronal networks may derive from patterns in neurovascular anatomy^29^: because neuronal and vascular growth processes track each other during development^30^, remote brain regions that establish neuronal links may also establish similar vascular, astrocytic, or other glial anatomy that influences local hemodynamic regulation. In the fully developed brain, environmental factors and repetitive activities (e.g., exercise) that impact the expression of neurotrophic factors may also simultaneously alter local angiogenesis^31–33^, allowing for ongoing and coregulated plasticity of neuronal and vascular networks. By coordinating blood flow across brain regions that typically exhibit synchronous neuronal activity, such vascular networks would also provide the most efficient hemodynamic support for increased network metabolism.

In addition to providing localized, responsive hemodynamic support for neuronal metabolism, the vasculature may also have a synergistic role in network brain activity: another interpretation of our findings is that vascular physiology *modulates* neuronal activity to drive the splitting of functional brain networks. Thus, the “more vascular” networks identified in this study may still fundamentally represent neuronal systems, but are somehow modulated by CO_2_ levels, whereas the associated “neuronal networks” are not specifically affected. There is emerging evidence that vascular physiology can influence neural activity^34–36^, and our lab has demonstrated that end-tidal CO_2_ changes, during gas inhalation and during resting fluctuations in breathing, can modulate neuronal rhythms as measured using magnetoencephalography (MEG)^37^. It has been further hypothesized that the vasculature may be directly involved in the brain’s information processing (the so-called hemo-neural hypothesis^38^), modulating the excitability of neural circuits via chemical, physical, and thermal mechanisms.

Importantly, these concepts are not mutually exclusive: there may be network-specific variation in vascular anatomy and regulation, neurovascular coupling, and vascular modulation of neural activity. It is not possible to differentiate these mechanisms in the current study. However, regardless of the precise origin of the observed relationships, we have demonstrated the dual nature of functional brain networks. It will be critical to ascertain how vascular physiology influences our interpretation of neuronal activity and connectivity within these systems.

Furthermore, our results support the recent work of Zhang and colleagues, who postulated the existence and importance of a “vascular-neural network” in understanding brain pathology^39^. This is an extension of the idea of the neurovascular unit, which has been a critical “conceptual framework” for understanding neurodegenerative disease and cerebrovascular injury^40^. The neurovascular unit includes endothelial cells, astrocytes, pericytes, and neurons, which must all interact in concert to maintain healthy neural function; the combined behavior of the unit must be considered when characterizing disease processes or developing new neuroprotective strategies^39^. However, the neurovascular unit spans less than a millimeter, and does not include upstream arteriolar supply vessels or downstream venous drainage. By linking these components, the vascular-neural network construct provides a useful integrated model that better describes systemic and focal neurovascular pathology, including in Alzheimer’s Disease, Parkinson’s Disease, multiple sclerosis, and autoimmune diseases of the central nervous system^39^.

At present, we may be missing key factors in disease progression by ignoring such long-range vascular systems. We know that early stages of ischemia affect both neurons and their supply microvessels in concert^40^. In Alzheimer’s Disease, vascular damage can precede and drive neurodegeneration^41,42^. Our results indicate that such pathological changes in neuro-vascular interactions could be network specific. This has been supported by recent research showing that healthy adults with vascular risk factors showed impairments in cerebrovascular reactivity to CO_2_ that were specific to the Default Mode Network, which is considered central to pathological mechanisms in aging and Alzheimer’s Disease^43^. Improving our understanding of vascular network behavior (or the vascular-neural network construct), and how the vasculature in specific functional networks is susceptible to early pathological impairment, may offer new windows for targeted protective therapies.

There is some existing evidence in the literature for pairs of spatially-similar networks observed in resting state fMRI data. Braga and Buckner identified two similar, but distinct, networks that both resembled the canonical Default Mode Network^44^. Other networks, including the dorsal attention network and fronto-parietal network were also fractioned into two distinct, parallel networks within individual datasets. The authors hypothesize that these are neuronal sub-networks, but the role of vascular physiology in network-specific BOLD signals was not considered. It may be that one or more of these observed sub-networks is primarily a vascular network, or that vascular regulation is altering BOLD signal fluctuations in specific subnetworks to drive their differentiation.

There are numerous challenges involved with using BOLD fMRI to study simultaneous neural and vascular properties of the brain. The BOLD contrast mechanism reflects local levels of deoxygenated hemoglobin, but this is constantly modulated by both direct vasoactive pathways and indirect neurovascular coupling mechanisms. Alternative imaging modalities such as EEG and MEG may provide better direct insight into the neural processes underpinning functional brain networks, however it is not yet fully understood how vascular physiology may manifest in these data^37^ or how network activity fluctuations in these different modalities relate back to fMRI signals^45^. In this study, the dual nature of BOLD fMRI contrast, in conjunction with dual stimulus types, facilitates our ability to probe the dual nature of functional brain networks, but perhaps future research should employ multi-modal imaging to best explore these phenomena in greater detail.

It is also challenging to employ spatial ICA to extract these spatially similar networks, as the algorithm maximizes the spatial independence of the resulting components. However, temporal ICA is not well suited to fMRI data, particularly when acquired using “typical” sampling rates of 1-2 seconds, due to the small number of degrees of freedom in each dataset. We also arbitrarily opted to decompose the data into 30 components; further assessment of other output dimensionality at this step in the analysis did impact the identification of network pairs, suggesting that the precise “splitting” of networks is highly dependent on this analysis choice. Similar observations have been observed by other research groups, where increasing ICA dimensionality facilitates the differentiation of sub-network structures^46^. Further studies into neuronal and vascular network properties should carefully assess the role of dimensionality on our observations, adopting rapid-sampling EPI (using simultaneous multi-slice acceleration to achieve sub-second sampling^47^) and testing the utility of temporal ICA at differentiating the neuronal and vascular features in the data.

## CONCLUSIONS

We have shown that functional brain networks can be split into two spatially similar networks during concurrent neuronal and vascular stimuli. One of these networks is dominated by the neuronal stimulus paradigm, as expected, whereas the other network appears dominated by vasodilatory responses to changes in arterial CO_2_ levels. This suggests that vascular regulation may be coordinated across long-distance brain regions, mimicking the structure of neuronal networks, or that neurovascular relationships vary in a network-specific manner. It will be critical to consider how the underlying vascular function influences the observation and interpretation of network brain activity and connectivity in future neuroimaging studies.

## Supporting information

## Acknowledgments

This work was supported by the Wellcome Trust [090199] and the Anne McLaren Fellowship program.

## Author Contributions

M.G.B. and K.M. designed this study. M.G.B., J.R.W., I.D.D., and K.M. acquired the fMRI data. M.G.B. analyzed the fMRI data. M.G.B. and K.M. prepared the manuscript. All authors discussed results and commented on the final manuscript.

