## Supplementary material for "Vascular physiology drives functional brain networks"

**Supplemental Figures:**

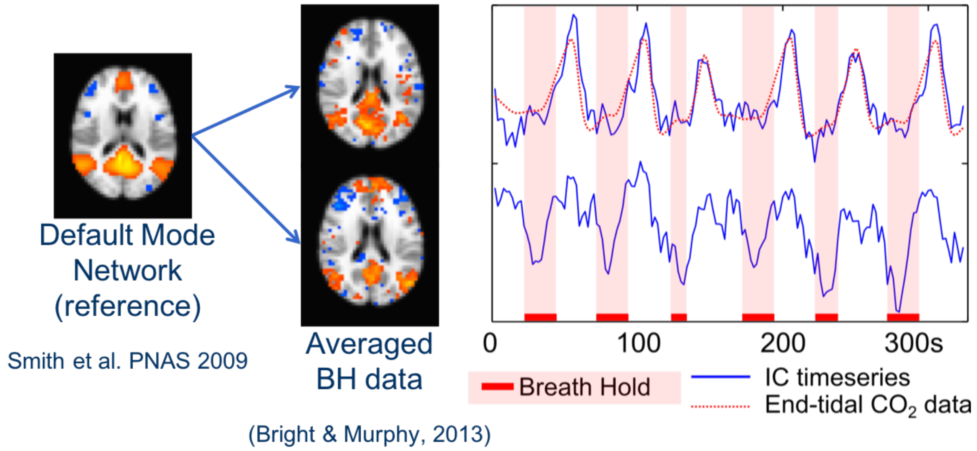


**Figure S1.** Decomposition of averaged breath-hold fMRI data using ICA resulted in the Default Mode Network being represented in two components. The component maps and associated time-series (blue) are presented, with the breath-hold timing and average end-tidal $CO_2$ trace (red). The top component shows BOLD signal increases closely following the end-tidal $CO_2$ trace. The bottom component shows BOLD signal decreases preceding and during the breath-hold stimulus, possibly reflecting deactivation of the neuronal Default Mode Network during the execution of the paced-breathing and breath-hold task. These results drove our hypothesis that functional brain networks, like the Default Mode Network, may be comprised of both neuronal and vascular systems.

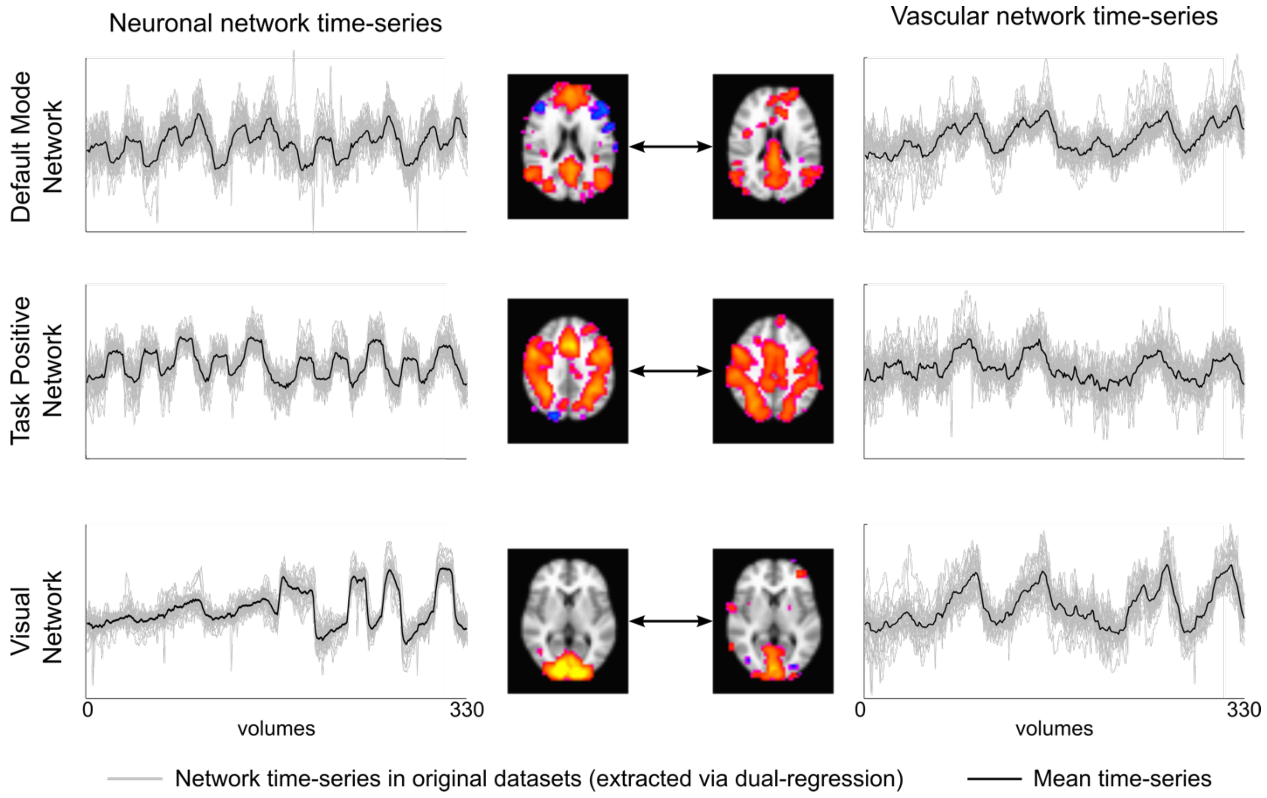


**Figure S2.** Time-series for vascular-neuronal network pairs. Time-series were extracted from the original 30 datasets using dual-regression. Networks of the left were identified via temporal correlation with the neural stimuli. Networks on the right were identified as having similar spatial characteristics to the neuronal network maps.
